# Genomic and phenotypic stability of fusion-driven pediatric Ewing sarcoma cell lines

**DOI:** 10.1101/2023.11.20.567802

**Authors:** Merve Kasan, Jana Siebenlist, Martin Sill, Rupert Öllinger, Enrique de Álava, Didier Surdez, Uta Dirksen, Ina Oehme, Katia Scotlandi, Olivier Delattre, Martina Müller-Nurasyid, Roland Rad, Konstantin Strauch, Thomas G. P. Grünewald, Florencia Cidre-Aranaz

**Affiliations:** Hopp Children’s Cancer Center (KiTZ), Heidelberg, Germany; Division of Translational Pediatric Sarcoma Research (B410), German Cancer Research Center (DKFZ), German Cancer Consortium (DKTK), Heidelberg, Germany; National Center for Tumor Diseases (NCT), NCT Heidelberg, a partnership between DKFZ and Heidelberg University Hospital, Heidelberg, Germany; Max-Eder Research Group for Pediatric Sarcoma Biology, Institute of Pathology, Faculty of Medicine, LMU Munich, Munich, Germany; Division of Pediatric Neurooncology, German Cancer Research Center (DKFZ); TranslaTUM, Center for Translational Cancer Research, Technical University of Munich, Munich, Germany; Institute of Biomedicine of Sevilla (IBiS), Virgen del Rocio University Hospital/CSIC/University of Sevilla/CIBERONC, Seville, Spain; Department of Normal and Pathological Cytology and Histology, School of Medicine, University of Seville, Seville, Spain; INSERM U830, Diversity and Plasticity of Childhood Tumors Lab, PSL Research University, SIREDO Oncology Center, Institut Curie Research Center, Paris, France; current address: Balgrist University Hospital, Faculty of Medicine, University of Zurich (UZH), Zurich, Switzerland; Department of Pediatrics, University Hospital Essen, Essen, Germany; Clinical Cooperation Unit Pediatric Oncology, German Cancer Research Center (DKFZ) and German Cancer Consortium (DKTK), 69120 Heidelberg, Germany; Experimental Oncology Laboratory, IRCCS Istituto Ortopedico Rizzoli, Bologna, Italy; Institute of Medical Biostatistics, Epidemiology and Informatics (IMBEI), University Medical Center, Johannes Gutenberg University, Mainz, Germany; IBE, Faculty of Medicine, LMU Munich, Munich, Germany; Institute of Genetic Epidemiology, Helmholtz Zentrum München - German Research Center for Environmental Health, Neuherberg, Germany; Department of Medicine II, Klinikum Rechts der Isar, Technical University Munich, Munich, Germany; German Cancer Consortium (DKTK), German Cancer Research Center (DKFZ), Munich, Germany; Institute of Pathology, Heidelberg University Hospital, Heidelberg, Germany

**Keywords:** Cancer models, cancer genomics, reproducibility, cell biology, sarcoma, carcinoma

## Abstract

Human cancer cell lines are the mainstay of cancer research. Recent reports showed that highly mutated adult carcinoma cell lines (mainly HeLa and MCF-7) present striking diversity across laboratories and that long-term continuous culturing results in genomic/transcriptomic heterogeneity with strong phenotypical implications. This highlighted how despite human cell line models being powerful tools for cancer research, the findings derived from their use may present limitations in terms of reproducibility. However, to what extent these conclusions can be generalized to the majority of cancer cell lines remained unexplored. Here, we hypothesized that oligomutated pediatric sarcoma cell lines driven by a chimeric oncogenic transcription factor (COTF), such as Ewing sarcoma (EwS), were genetically and phenotypically more stable than the previously investigated (adult) carcinoma cell lines. A comprehensive molecular and phenotypic characterization of multiple EwS cell line strains in direct comparison to the HeLa and MCF-7 cell lines, together with a simultaneous analysis during 12 months of continuous cell culture showed that COTF-driven pediatric sarcoma cell line strains are genomically more stable than adult carcinoma strains, display remarkably stable and homogenous transcriptomes, and exhibit uniform and stable drug response. The analysis of multiple EwS cell lines subjected to long-term continuous culture conditions revealed that variable degrees of genomic/transcriptomic/phenotypic may be observed among COTF-driven cell lines, further exemplifying that the potential for reproducibility of in vitro scientific results may be rather understood as a spectrum, even within the same tumor entity.

## MAIN TEXT

Cancer cell lines have been instrumental for biomedical progress for many decades^1–3^. In 2018 and 2019, respectively, Ben-David *et al.* and Liu *et al.* showed that highly mutated adult carcinoma cell lines present striking diversity across laboratories and that long-term continuous culturing results in genomic/transcriptomic heterogeneity with phenotypical implications, including changes in drug sensitivity^4^, doubling time, and response to a specific perturbation^5^, which challenged the general reproducibility of scientific results based on human cancer cell lines. However, to which extent these observations can be generalized to every cancer cell line remains to be explored. Here, we hypothesized that oligomutated pediatric sarcoma cell lines driven by a chimeric oncogenic transcription factor (COTF), such as Ewing sarcoma (EwS)^6^ are genetically and phenotypically more stable than the previously investigated (adult) carcinoma cell lines^4,5^.

The multi-omics study by Liu *et al.* showed a substantial heterogeneity between different variants of the first human-derived cancer cell line HeLa (cervix carcinoma)^7^, mainly between the most commonly used variants HeLa-CCL2 and HeLa-Kyoto. Interestingly, Ben-David *et al.* reanalyzed the genomic data (whole exome sequencing) of 106 cancer cell lines provided by the Broad and the Sanger Institutes and showed a significant diversity in allelic fraction for somatic variants in this panel of cell lines. Notably, this panel mainly consisted of hematopoietic/lymphoid and adult carcinoma cell lines and only included a single EwS cell line (CADO-ES1), which was not further investigated. Among those adult carcinoma cell lines, the authors specifically focused on the estrogen receptor (ER)-positive adult breast carcinoma cell line MCF-7 for a cross-laboratory analysis and demonstrated a crucial genomic, transcriptomic, and phenotypical diversity. They additionally verified their findings in a panel of adult carcinoma cell lines including (except for a single pediatric hepatoblastoma cell line HepG2-A) mostly adult-type carcinoma cell lines, all of which are not driven by a single mutation, such as the COTFs found in EwS.

Hence, to test whether COTF-driven sarcoma cell lines are clonal or genetically unstable, we selected human A-673, one of the most widely used EwS cell line, and compared 11 A-673 strains with five strains of human HeLa cervix cancer and five strains of human MCF-7 breast cancer collected from seven, three, and two different laboratories, respectively (**Fig. 1a**). Despite some of these strains had an undefined number of passages, they were considered serviceable for cell biology research. In this comparison, we included a newly purchased strain for each cell line, which were continuously cultured for 12 months, and examined at three different time points (corresponding to month 0, 6, and 12, hereafter referred to as m0, m6, m12) (**Fig. 1a**). To reduce empirical bias prior to our (epi)genomic, transcriptomic, and phenotypical analyses, we cultured all strains in the same cell culture conditions (see Methods section).

**Fig. 1.**
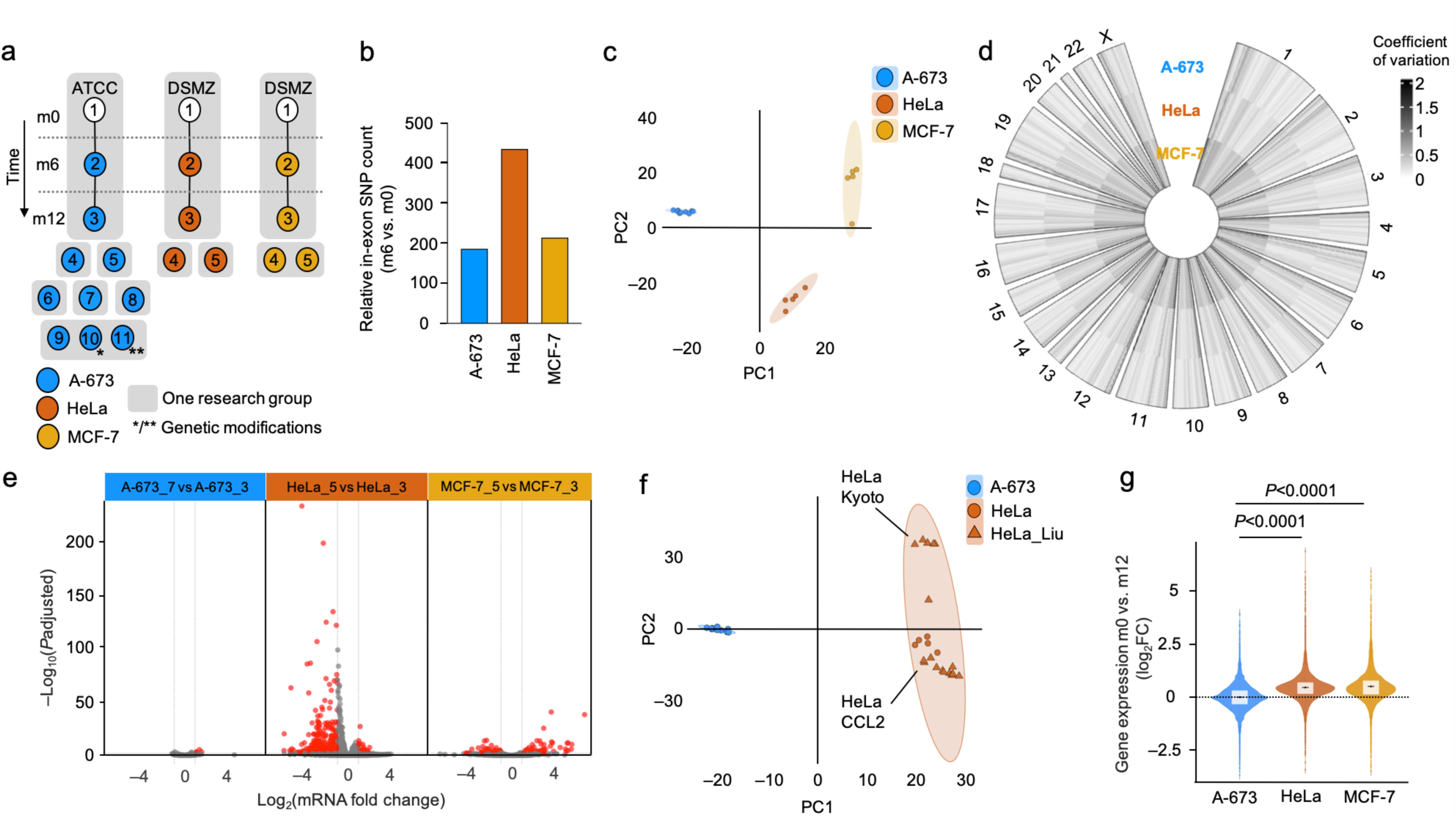
COTF-driven pediatric sarcoma cell line strains exhibit exceptional genomic and transcriptomic stability compared to adult carcinoma strains. **a.** Newly purchased A-673 EwS, HeLa cervix carcinoma and MCF-7 breast carcinoma wild type cell lines (A-673_1, HeLa_1 and MCF-7_1) were kept in culture for six months (m6; A-673_2, HeLa_2 and MCF-7_2) and twelve months (m12; A-673_3, HeLa_3 and MCF-7_3). Additionally, multiple strains for each cell line were collected from seven, three and two laboratories, respectively, and arbitrarily numbered as A-673_4 to A-673_9, HeLa_4 to HeLa_5, and MCF-7_4 to MCF-7_5. Single cell clones derived from A-673 cell lines with a neutral manipulation (*, A-673_10) or an inducible shRNA construct targeting its EWSR1::FLI1 translocation (**, A-673_11) were included. ATCC, American Type Culture Collection, DSMZ (German Collection of Microorganism and Cell Cultures). **b.** Relative in-exon SNP counts for A-673, HeLa, and MCF-7 after six months of continuous culture, using the respective initial time point (m0) values as reference (m6 vs. m0). **c.** Transcriptomic PCA of 11 A-673, five HeLa and five MCF-7 cell line strains (*N*=10,256 transcripts). **d.** Circle plot depicting coefficient of variation (CV) of expressed genes per chromosome (top 60% quantile) for all A-673, HeLa and MCF-7 cell line strains. **e.** Volcano plot of DEG resulting from comparing the two A-673, HeLa and MCF-7 strains with the highest variance (A-673_7 vs A-673_3, HeLa_5 vs HeLa_3 and MCF-7_5 vs MCF-7_3). The red dots denote significantly differentially expressed genes (BH adjusted *P*<0.01; |FC|>1). **f.** Combined transcriptomic PCA of our A-673 and HeLa dataset and that of Liu *et al.* (HeLa_Liu) (*N*=13,569 transcripts) **g.** Relative gene expression of A-673, HeLa, and MCF-7 cell lines after long-term culture for twelve months (m12 vs m0). Outer violin curves denote the kernel density. Boxplots display the interquartile range and the mean, two-sided Wilcoxon signed-rank test.

In a first step, we performed a cross-strain analysis of A-673, MCF-7, and HeLa cell lines and genotyped each newly purchased cell line (m0) and its respective m6-cultured version using Illumina Global Screening Arrays (GSA), which enabled us to monitor genetic evolution over time at the level of point mutations at reference SNPs represented on this array type. In keeping with the consistent results of cell line authentication using STR profiling, over 98% of the genotyped SNPs were identical for each given cell line after six months of continuous culture. However, when analyzing in-exon SNP counts, we noted genetic evolution over time that was 2.33-fold and 1.15-fold higher in HeLa and MCF-7 cells, respectively, compared to A-673 EwS cells (**Fig. 1b**). Further, analysis of non-synonymous SNPs that affected the coding sequence and splicing regions for the 11 different A-673 strains (including two A-673 with genetic modifications) revealed that 98.9% were shared by all strains (**Supp. Fig. 1a**), which drastically diverged from the only 35% of SNPs shared by all strains in MCF-7^4^.

To investigate this discrepancy between adult carcinoma cell lines and oligomutated pediatric sarcomas at the transcriptional level, we compared the transcriptomic variation of these previously studied adult carcinoma cell lines with COTF-driven EwS cell lines. Specifically, we performed RNA sequencing (RNASeq) using NextSeq 500 (Illumina) on 11 A-673, five HeLa, and five MCF-7 strains. Principal component analysis (PCA) performed on the transcriptomic data, from three biological replicates per cell line, revealed that similar to the observations made by Ben-David *et al*. and Liu *et al.*^4,5^ the strains of both carcinoma cell lines showed widespread transcriptomic diversity. However, our COTF-driven A-673 EwS strains clustered tighter than HeLa and MCF-7 carcinoma strains, even despite the A-673 cluster contained two strains with genetic modifications, and both carcinoma cell lines had relatively smaller sample sizes (**Fig. 1c**). These observations were additionally confirmed by analyzing the coefficient of variation (CV) of gene expression for each cell line (**Fig. 1d**). To study this phenomenon in more detail, we compared for each cancer entity the two strains with the highest variance (A-673_7 and A-673_3 vs. HeLa_5 and HeLa_3 vs MCF-7_5 and MCF-7_3). Strikingly, we observed over 60 times more differentially expressed genes (DEG) defined as |fold change (FC)|>1, Benjamini-Hochberg (BH) adjusted *P*<0.01 (380 transcripts; 39 up-, 341 down-regulated) in the HeLa strains and 20 times more DEG (108 transcripts; 57 up-, 51 down-regulated) in the MCF-7 strains as compared to the A-673 EwS strains (5 transcripts; all up-regulated) (**Fig. 1e**, **Supp. Fig. 1b**). We additionally combined our transcriptomic data with that of Liu *et al.* and observed a specific clustering of our HeLa strains with their HeLa-CCL2 strains, indicating a likely common origin (**Fig. 1f**). Moreover, we observed a remarkably higher degree of heterogeneity among HeLa strains than among A-673 strains (**Fig. 1f**, **Supp. Fig. 1c**). Of note, considering this heterogeneity between HeLa-CCL2 and Kyoto strains described by Liu *et al.* and here, it is conceivable that the inclusion of HeLa-Kyoto in our panel of cells (**Fig. 1a**) would have resulted in an even more substantial difference when compared to COTF-driven A-673.

Next, we compared the expression profiles of the newly purchased cell lines (m0) for each cancer entity with their m12 derivates. Consistent with the results observed in the cross-laboratory comparison, we observed a significantly greater variation in global gene expression (*P*<0.0001, two-sided Wilcoxon signed-rank test) in HeLa and MCF-7 cells compared to A-673 (median log_2_FC_A-673_=0, –4.25<X̃<4.22; median log_2_FC_HeLa_=0.47, –3.39<X̃<16.89; median log_2_FC_MCF-7_=0.47, –3.46<X̃<15.81) (**Fig 1g**, **Supp. Fig. 1d**).

To evaluate the potential phenotypical impact of these genomic and transcriptomic changes, we compared the drug responses of COTF-driven EwS cells with those from highly mutated adult carcinomas. Thus, we subjected 11 A-673 EwS strains (including two A-673 with genetic modifications, **Fig. 1a**), five HeLa cervical cancer strains, and five MCF-7 breast cancer strains to a drug screening consisting of a selection of 10 active compounds addressing non-redundant functional pathways, which encompassed the same drugs used in Ben-David *et al.*^4^. The obtained dose-response curves were used to compute the area under the curve (AUC) for each compound and to determine the respective Euclidean distances (ED) between sensitivity profiles of a given cell line to the global AUC-mean across cell lines. In agreement with our previous findings, the strains of both adult carcinomas exhibited a significantly higher degree of drug response variability than those of EwS for all screened compounds (**Fig. 2a**, *P*<0.005, one-sided Wilcoxon signed-rank test). To confirm the extensive homogeneity in drug response of the COTF-driven EwS cells as compared to carcinoma cell lines, we additionally performed a Spearman’s correlation test among each cell line’s strains and once again observed that EwS strains showed a higher similarity than carcinoma cell lines (X̃_Spearman’s *ρ* A-673_=0.94, 0.95<X̃<0.93; X̃_Spearman’s *ρ* HeLa_=0.87, 0.91<X̃<0.83; X̃_Spearman’s *ρ* MCF-7_=0.88, 0.92<X̃<0.85) (**Fig. 2b**).

**Fig. 2.**
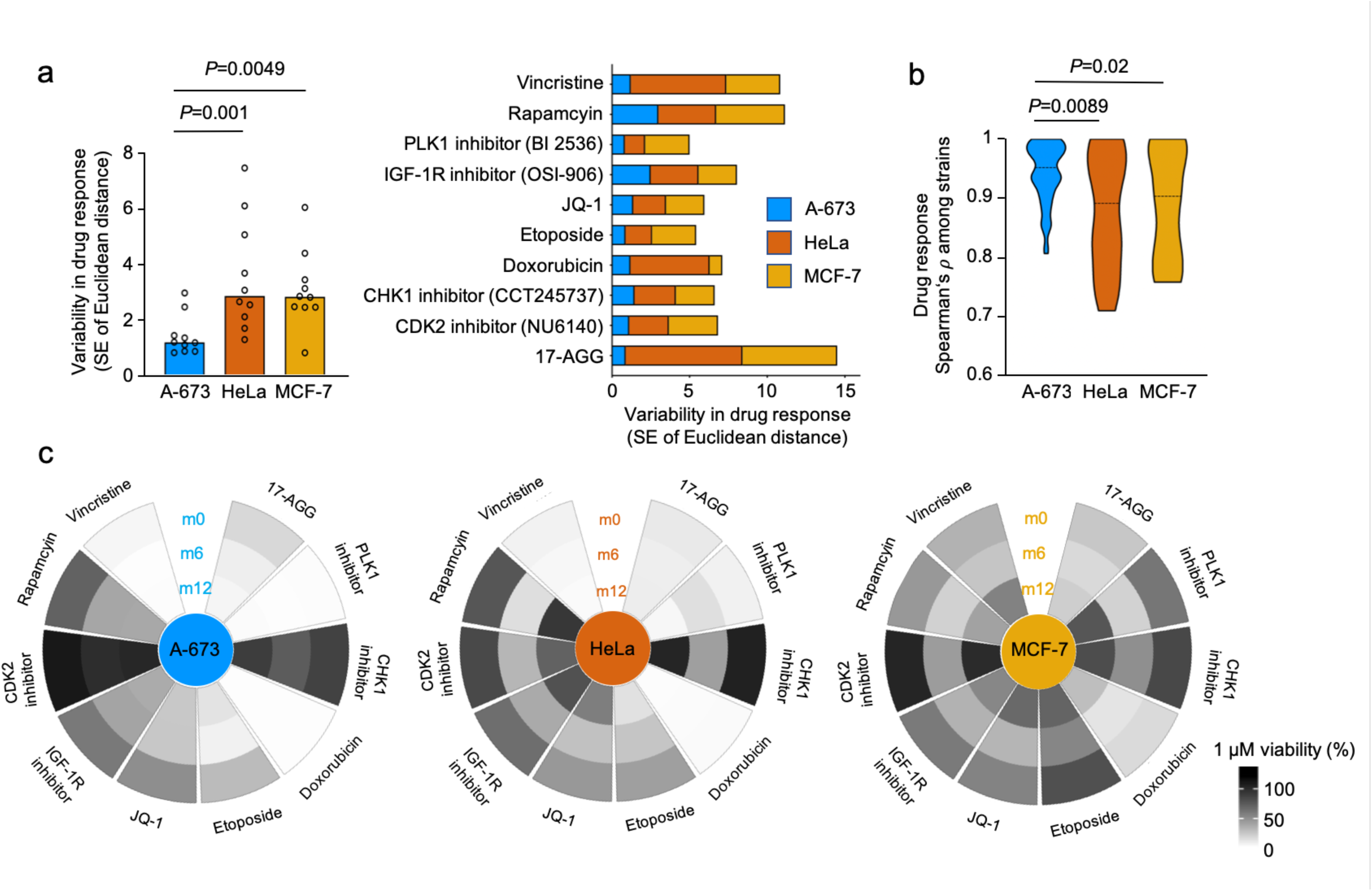
Uniform and stable drug-response in COTF-driven pediatric sarcoma cell lines. **a.** Left, collective variability in drug response of all A-673, HeLa, and MCF-7 cell line strains depicted as standard error of ED, each compound is shown as a black circle, one-sided Wilcoxon signed-rank test. Right, standard error of ED for each specific screened compound. **b.** Spearman’s *ρ* of drug response across 11 A-673, five HeLa, and five MCF-7 cell line strains. Dotted black line shows the median (one-sided Wilcoxon rank-sum test). **c.** Raw viability of A-673, HeLa and MCF-7 cell lines subjected to each compound (1 µM) included in the drug screening after 0, 6, and 12 months of continuous long-term culture (m0, m6, and m12).

Further, we studied the effect of continuous culture over 12 months on the potential evolution in drug sensitivity. Therefore, we exposed newly purchased A-673, MCF-7, and HeLa cells (m0) to the drug library, and then again at two additionally predefined time points after continuous culture (m6 and m12). In agreement with previous findings, we detected a remarkably stable phenotype of A-673 after 6 and 12 months in comparison with HeLa and MCF-7 cell lines, measured as raw viability at a single concentration of each compound (1 µM) (**Fig. 2c**).

Finally, to expand our understanding of the scarcity of genomic and phenotypic cell line evolution in the context of EwS, we sought to analyze how our findings in A-673 cells (as one of the most widely used cell line in EwS research) would compare to other EwS cell lines. Thus, we newly purchased four additional EwS cell lines (MHH-ES1_m0, SK-ES-1_m0, SK-N-MC_m0 and TC-71_m0) and propagated them for 12 months (**Fig. 3a**). We first performed genomic and epigenomic analyses and subjected our samples to Illumina GSA and MethylationEPIC BeadChip arrays, respectively. Interestingly, while all EwS cell lines remained relatively stable, we observed that when compared to the prototypical A-673 cell line, the remaining EwS cell lines presented a gradient of variability when analyzing both their non-synonymous SNP alterations and their differentially methylated CpG sites over time (**Fig. 3b,c**). For instance, while a median of 99.6% (range 99.3%-99.8%) of the in-exon SNPs were shared after 12 months of continuous culture by each cell line, we could observe relatively less stable cell lines such as A-673, and MHH-ES-1, and remarkably stable cell lines such as TC-71 (**Fig. 3b**), whereas SK-ES-1 and A-673 displayed less stability at epigenetic level (**Fig. 3c**). This observed genomic and epigenomic variability in degree of evolution over time was further detected at the transcriptional level, as shown by the proportion of significantly DEG of each EwS cell line when compared to their m12 derivate (**Fig. 3d**). Indeed, TC-71 showed the least transcriptional changes over time, while SK-ES-1 exhibited the highest number of DEG after 12 months of continuous culture (219 transcripts; 99 up- and 120 down-regulated, which represented a 50% increment relative to A-673) (**Fig. 3d**).

**Fig. 3.**
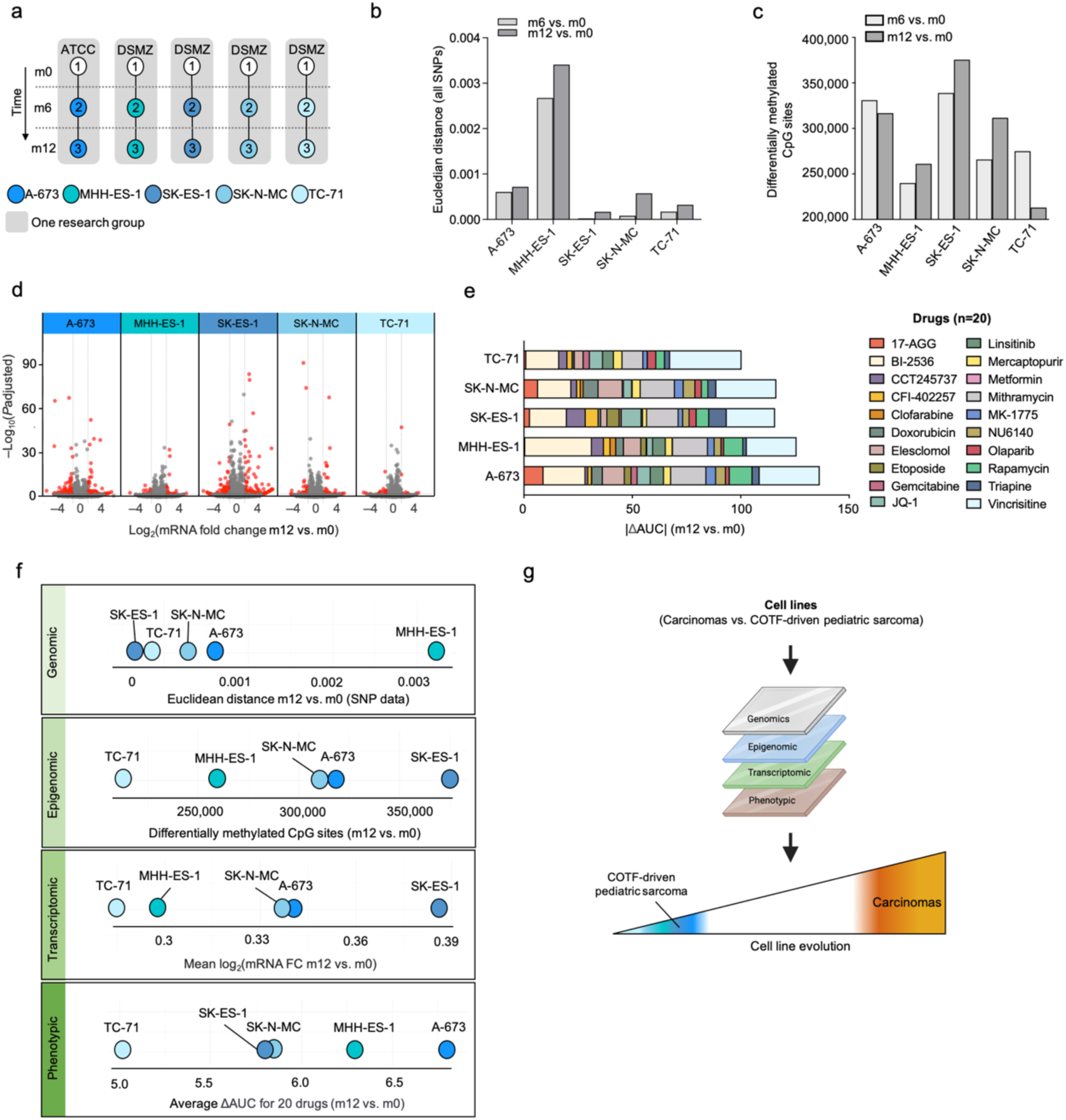
In-depth analysis of stability on individual cell lines from the same COTF-driven sarcoma entity. **a.** Newly purchased A-673, MHH-ES-1, SK-ES-1, SK-N-MC, and TC-71 EwS wild type cell lines (_1) were kept in culture for six months (m6; _2), and 12 months (m12; _3). ATCC, American Type Culture Collection, DSMZ (German Collection of Microorganism and Cell Cultures). **b.** Bar plot shows the Euclidean distance of all SNPs after 6 (m6 vs. m0) and 12 months (m12 vs. m0) of continuous culture. **c.** Number of differentially methylated CpG sites (including differentially hypo-and hyper-methylated) for A-673, MHH-ES-1, SK-ES-1, SK-N-MC, and TC-71 after six (m6) and 12 months (m12) of continuous culture, using the respective initial time point (m0) values as reference. **d.** Volcano plot of DEG comparing each EwS cell line with their m12 derivate. Red dots denote significant DEG (BH adjusted *P*<0.01; |FC|>1). **e.** Relative variation in cell viability for each EwS cell line measured as mean area under the curve (AUC) m12 vs. m0 for the extended drug library (20 compounds). **f.** Ranking plots depicting the linear distribution of five EwS cell lines based on their evolution across different datasets after continuous culturing for 12 months. **g.** Schematic illustration summarizing the findings of this study.

We next complemented these results by exposing each newly purchased EwS cell line (m0) and their 12-month derivate (m12) to an extended drug library that contained 10 additional compounds (extended library, n_total_=20, **Supp. Table 1**) to include drugs that had been recently described in EwS preclinical or clinical studies, such elesclomol, olaparib and gemcitabine^8–10^. Here, we again observed inter-cell line variability in collective drug response over time that ranged from the least stable A-673, to the remarkably stable TC-71 EwS cell line (**Fig 3e**). In synopsis, ranking plots for each different data layer comparing 12 months of continuous culture of each EwS cell line clearly suggest a range of stability that may inform decision-making on which cell line models to preferentially employ in this COTF-driven pediatric cancer (**Fig. 3f**).

Collectively, our results widely confirm those of Liu *et al.* and Ben-David *et al*., while highlighting that their findings may not be translatable to other cell lines, especially to those with a stable genetic background and a defined driver mutation, such as the COTF found in EwS (**Fig. 3g**). Our findings indicate that research with COTF-driven cell line models such as EwS should be in principle reproducible, even after genetic modifications, and extensive periods of continuous culture. Also, our results demonstrate that individual cell lines from the same cancer entity may display a variable degree of evolution, suggesting that the reproducibility of results strongly depends on the given cancer cell lines.

## MATERIALS AND METHODS

### Provenience of cell lines and cell culture conditions

For long-term culture assays the following early passage (<5 passages) human cancer cell lines were acquired: the cervix carcinoma cell line HeLa, the human breast carcinoma cell line MCF-7. The EwS MHH-ES-1, SK-ES-1, SK-N-MC, and TC-71 cell lines were purchased from the German Collection of Microorganism and Cell Cultures (DSMZ). The A-673 EwS cell line was purchased from the American Type Culture Collection (ATCC). A-673, HeLa, and MCF-7 wild-type strains with an undefined number of passages were collected from different laboratories. Single cell clones derived from A-673 cell lines with either a neutral manipulation (A-673/shcontrol) or an inducible shRNA construct against its EWSR1::FLI1 translocation (A-673/TR/shEF1) were previously described by our laboratory^11^. All cell lines were routinely tested for mycoplasma contamination by nested PCR, and cell line purity and authenticity were confirmed by STR-profiling. All cell lines were cultured at 37 °C, 5% CO_2_ in RPMI 1640 (Biochrom, Germany) supplemented with 10% fetal bovine serum (Sigma-Aldrich, Germany) and 1% penicillin-streptomycin (Merck, Germany). Each cell culture flask was monitored daily and cells were passaged twice per week using Trypsin-EDTA (0.25%) (Life Technologies) when they reached approximately 70% confluency.

### DNA extraction, methylation and global screening arrays

When flasks reached approximately 70% confluency, samples were lysed, and total DNA was extracted with the NucleoSpin Tissue kit (Macherey Nagel) following the manufacturer’s protocol. For each sample, 900 ng of DNA in one (genoryping) or two (methylation) technical replicates were used as input material and were profiled on Illumina Infinium Global Screening array and MethylationEPIC array, respectively, at the Molecular Epidemiology Unit of the German Research Center for Environmental Health (Helmholtz Center, Munich, Germany).

### DNA methylation data analysis

The initial pre-processing of the raw methylation was performed in R version 3.3.1. Raw signal intensities were obtained from IDAT-files using the minfi Bioconductor package version 1.21.4 (ref^12^) in R version 3.3.1. Each sample was individually normalized by performing a background correction (shifting of the 5% percentile of negative control probe intensities to 0) and a dye-bias correction (scaling the mean of normalization control probe intensities to 10,000) for both color channels. The methylated and unmethylated signals were corrected individually. Subsequently, beta values were calculated from the retransformed intensities using an offset of 100 (as recommended by Illumina). Out of 865,859 probes on the EPIC array, 105,454 probes were masked according to Zhou et al.^13^ as well as 16,944 probes on the X and Y chromosomes. In total, 743,461 probes were kept for downstream analysis. The beta values were transformed to M-values with the logit2 function of the *minfi* package version 1.42.0, R version 4.2.0. A probe-wise differential methylation analysis^14^ was performed using the *limma* package^15^ version 3.52.4 in R version 4.2.0 by comparing six and twelve months of culturing with the initial time point (m0) as reference. Significant differentially methylated CpG probes were extracted with the *decideTests* function of the *limma* package with an FDR<0.05 (Benjamini-Hochberg). All significantly differentially methylated (total hypo- and hyper-methylated) CpG sites were visualized using PRISM 9 (GraphPad Software Inc. CA, USA).

### Global screening array (GSA) data analysis

The initial processing and quality control (QC) of the raw genotyping data was performed using PLINK version 1.9 (SNP call rate >95%, Hardy-Weinberg exact test <1e-6, and variants on Y chromosome were excluded). In total 526,610 variants out of 696,726 passed the QC filters. Infinium GSA v3.0 annotation file was used to filter for in-exon or non-synonymous variants. To determine single nucleotide alterations (SNA) after six and 12 months in cell culture (m6 and m12), each cell line was compared to its m0 version (number of consistent alleles and changes from homozygous to heterozygous) using the Variant Call Format (VCF) file generated by PLINK 1.9. Further data analysis was performed in R version 4.2.1, using the vcfR package version 1.14.0, among other data processing packages described below. The distance between two time points for each cell line was computed in R version 4.2.1 using the proxy package version 0.4-27. The eigenvectors, generated for dimension reduction in PLINK version 1.9, were used as input. The heatmap was generated in R version 4.2.1 using pheatmap package version 1.0.12.

### RNA extraction, library preparation, RNA sequencing and analysis

When flasks reached approximately 70% confluency, total RNA was isolated using the NucleoSpin RNA kit (Macherey-Nagel, Germany) according to the manufacturer’s protocol. RNA quality was verified on a Nanodrop Spectrophotometer ND-1000 (Thermo Fischer) and quantity was measured on a Qubit instrument (Life Technologies). For each sample, 50–100 ng of RNA in three biological and two technical replicates were used as input material and were profiled on an Illumina NextSeq 500 system at the Institute of Molecular Oncology and Functional Genomics in Rechts der Isar University Hospital (TranslaTUM Cancer Center, Munich, Germany). Library preparation for bulk 3’-sequencing of poly(A)-RNA was performed as previously described^16^. Briefly, barcoded cDNA of each sample was generated with a Maxima RT polymerase (Thermo Fisher) using oligo-dT primer containing barcodes, unique molecular identifiers (UMIs) and an adapter. 5’ ends of the cDNAs were extended by a template switch oligo (TSO) and after pooling of all samples full-length cDNA was amplified with primers binding to the TSO-site and the adapter. cDNA was fragmented and TruSeq-Adapters ligated with the NEBNext® Ultra™ II FS DNA Library Prep Kit for Illumina® (NEB) and 3’-end-fragments were finally amplified using primers with Illumina P5 and P7 overhangs. P5 and P7 sites were exchanged to allow sequencing of the cDNA in read1 and barcodes and UMIs in read2 to achieve better cluster recognition. The library was sequenced with 75 cycles for the cDNA in read1 and 16 cycles for the barcodes and UMIs in read2. Data was processed using the published Drop-seq pipeline (v1.0) to generate sample- and gene-wise UMI tables^17^. After elimination of transcripts with very low counts (sums of all samples <10), RNASeq data in count matrix format was batch corrected using ComBat-Seq function of R package sva version 3.44.0 (ref. ^18^), and differential gene expression analysis (DGEA) was performed using DESeq2 version 1.36.0 (ref. ^19^) on R version 4.2.1. Combat-Seq adjusted data was used as count input for DESeqDataSet. To avoid false discovery artifacts due to the detection of minimally expressed genes, we excluded the 40% lowest expressed genes across samples (remaining expressed genes *N*=10,257). For the analysis of long-term cultured EwS cell lines we performed DGEA on the top 60% expressed genes included in the raw count matrix (*N=*27,143, all EwS cell line samples were analyzed in one batch). For DGEA between two samples, genes with *P*_adj_≤0.01, |log2(FC)|>1 were considered as DEG. Principal component analysis was used to preserve the global properties of the data using the *plotPCA* function. To comprehensively display the degree of variability between strains in each tumor type, gene-specific CV of the transcriptomic data was calculated. In the long-term culture assays, log_2_FC of gene expression of each cell line for 6 and 12 months (m6 and m12) were analyzed using the initial time point (m0) values as reference.

### Drug screening

All A-673, HeLa and MCF-7 strains, as well as MHH-ES-1, SK-ES-1, SK-N-MC, and TC-71 EwS cell lines, were tested against a core drug library consisting of 10 cytotoxic or cytostatic agents, or an extended drug library consisting of 20 agents **(Supp. Table 1)**. For this, cells were seeded into 96-well plates at a density of 5×10^3^ cells per well in 90 µl of medium in triplicates. Once cells were attached, ~4 h after seeding, 10 µl of each compound was added in serially diluted concentrations ranging from 1×10^-5^ µM to 10 µM. DMSO was used as vehicle control. Plates were incubated for 72 h at 37 °C, with 5% CO_2_ in a humidified atmosphere. At the experimental endpoint, a solution of 25 µg/ml of resazurin salt (Sigma-Aldrich) was added to the medium and cell viability was determined as previously described^20^. Each compound and cell line were assayed in four biological replicates.

### Drug screening data analysis

Cell viability data was first normalized using the measured raw viability of each control (DMSO vehicle), and the area under curve (AUC) was computed for each cell line using the PharmacoGx package version 3.0.2 (P Smirnov, 2016) in R version 4.2.1. Euclidean distances (ED) between drug sensitivity profiles of each strain were calculated using the following formula: *function(x1, x2) sqrt(sum((x1* – *x2)*^2^*))= ED,* where x1 is the mean value of AUC of all strains and x2 the AUC of individual cell lines. The variability in drug response across different cancer entities was visualized using the standard error of ED values, accounting for differences in sample size. Changes in drug sensitivity during the long-term culture of each cell line for six and 12 months (m6 and m12) were analyzed using the initial time point (m0) values as reference.

### Other bioinformatic and statistical analyses

If not otherwise specified, genomic, methylation, transcriptomic, and drug sensitivity data analyses were performed in R version 4.2.1. The following R packages were used: for data processing, readxl package version 1.4.3, tidyverse package version 2.0. (ref.^21^), reshape2 package version 1.4.4 (ref.^22^), cowplot package version 1.1.1, Rfast package version 2.0.8, and data.table package version 1.14.8 (ref.^23^); for data visualization, ggplot2 package version 3.4.1 (ref.^24^), gghalves package version 0.1.4, ggdist package version 3.2.1 and PupillometryR package version 0.0.4; for circle plots, circlize package version 0.4.15; and for PCA and volcano plots, ggplot2 package version 3.4.1 (ref.^24^). Spearman’s correlation analyses of quantitative data of both mRNA and drug response were performed using Hmisc package version 4.7-2 (ref.^25^). Figures 1b, 2a, 2b, 3b, 3c, 3e, and Supplementary Figure 1c were generated using PRISM 9 (GraphPad Software Inc., Ca, USA). Transcriptomic datasets from this study and Liu *et al*.^5^ were combined and batch corrected using ComBat-Seq function of package sva version 3.44.0 (ref.^26^).

## DECLARATIONS

### Ethics approval and consent to participate

Not applicable.

### Data availability statement

Original data that support the findings of this study will be deposited at the National Center for Biotechnology Information (NCBI) GEO and will be accessible upon reasonable request.

### Code availability statement

Custom code is available from the corresponding author upon reasonable request.

## Acknowledgements

We thank Prof. Heinrich Kovar for kindly sharing materials, and Dr. Soledad Gómez-Gonzalez for critical discussion of this manuscript.

## Authors’ contributions

F.C.A. and T.G.P.G. conceived the study. M.K. performed all experiments, and bioinformatic and statistical analyses. F.C.A. contributed to drug screening experiments. J.S. contributed to bioinformatic analyses. M.S., R.Ö., R.R., M.M-N., and K.St. contributed to sample analysis and/or provided laboratory infrastructure. E.deÁ., D.S., O.D., U.D., I.Ö., and K.Sc. provided cell line models. M.K., F.C.A., and T.G.P.G. wrote the paper, and drafted the figures and tables. F.C.A and T.G.P.G. supervised the study and data analysis. All authors read and approved the final manuscript.

## SUPPLEMENTARY FIGURE LEGENDS

**Supp. Fig. 1.**
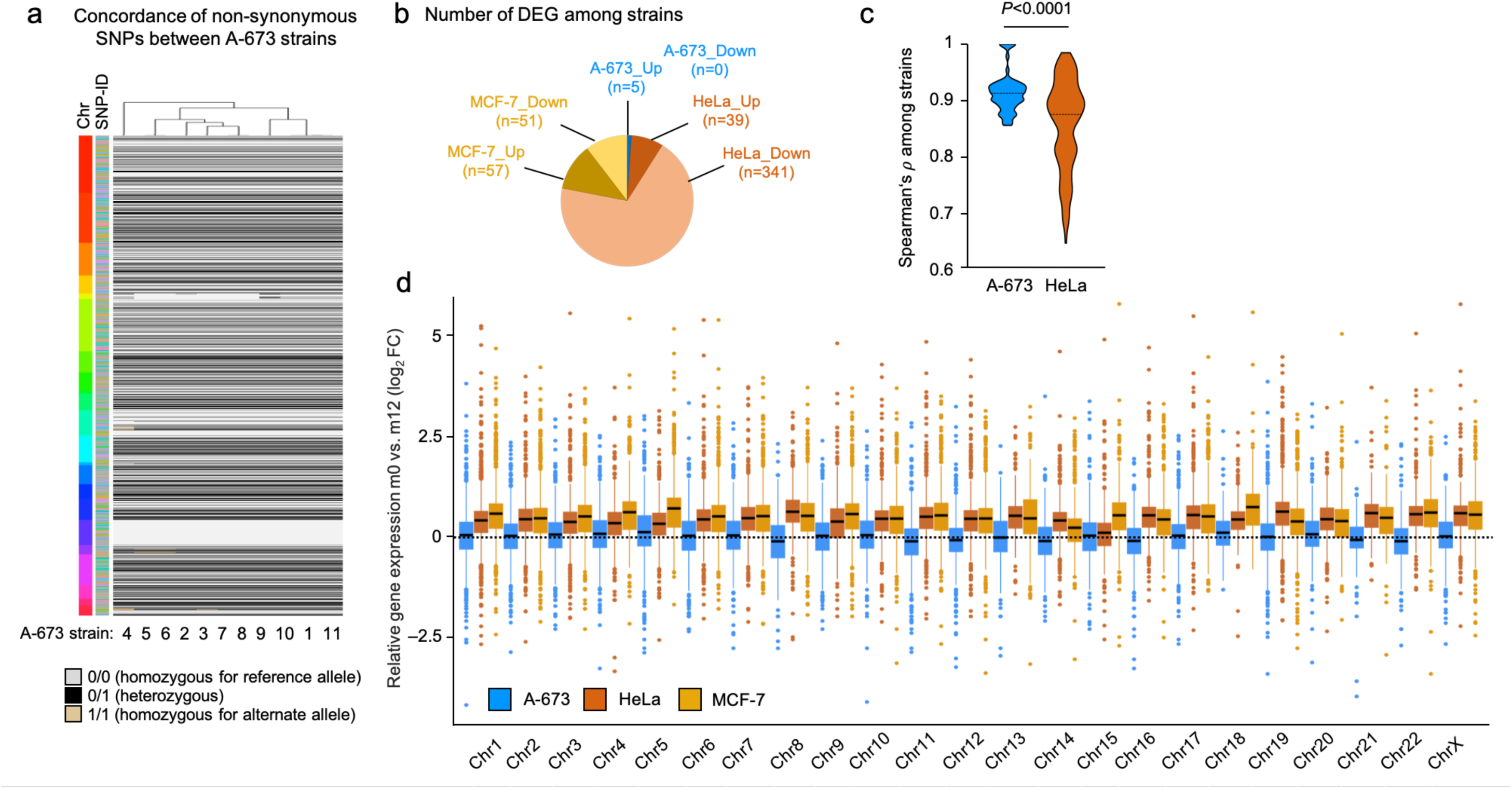
COTF-driven pediatric sarcoma cell line strains display stable and homogenous genomes and transcriptomes. **a.** Heatmap comparing the status (homozygous for reference allele, alternate allele, or heterozygous) of non-synonymous SNPs in 11 A-673 strains. Left color bar depicts chromosomes; right color bar depicts different SNP-IDs (*N*=1,599). **b.** Pie chart depicts the number of up-and down-regulated DEG emerging from each comparison. **c.** Gene-specific Spearman’s *ρ* across A-673 and HeLa cell line strains from the combined transcriptomic datasets in Fig 1f. Dotted black lines show the median (one-sided Wilcoxon rank-sum test). **d.** Relative gene expression of A-673, HeLa, and MCF-7 cell lines after long-term culture for twelve months (m12 vs m0) per chromosome. Boxplots display the interquartile range and the mean.

## REFERENCES

1. Iorio, F. et al. A Landscape of Pharmacogenomic Interactions in Cancer. Cell 166, 740–754 (2016).

2. Barretina, J. et al. The Cancer Cell Line Encyclopedia enables predictive modelling of anticancer drug sensitivity. Nature 483, 603–607 (2012).

3. Gonçalves, E. et al. Pan-cancer proteomic map of 949 human cell lines. Cancer Cell 40, 835–849.e8 (2022).

4. Ben-David, U. et al. Genetic and transcriptional evolution alters cancer cell line drug response. Nature 560, 325–330 (2018).

5. Liu, Y. et al. Multi-omic measurements of heterogeneity in HeLa cells across laboratories. Nat Biotechnol 37, 314–322 (2019).

6. Grünewald, T. G. P. et al. Ewing sarcoma. Nat Rev Dis Primers 4, 5 (2018).

7. Gey G.O, Coffman W.D., & Kubicek M.T. Tissue culture studies of the proliferative capacity of cervical carcinoma and normal epithelium. Cancer Res. 12, 264–265 (1952).

8. Marchetto, A. et al. Oncogenic hijacking of a developmental transcription factor evokes vulnerability toward oxidative stress in Ewing sarcoma. Nat Commun 11, 2423 (2020).

9. Takagi, M. et al. First phase 1 clinical study of olaparib in pediatric patients with refractory solid tumors. Cancer 128, 2949–2957 (2022).

10. Mora, J. et al. GEIS-21: a multicentric phase II study of intensive chemotherapy including gemcitabine and docetaxel for the treatment of Ewing sarcoma of children and adults: a report from the Spanish sarcoma group (GEIS). Br J Cancer 117, 767–774 (2017).

11. Orth, M. F. et al. Systematic multi-omics cell line profiling uncovers principles of Ewing sarcoma fusion oncogene-mediated gene regulation. Cell Rep 41, 111761 (2022).

12. Aryee, M. J. et al. Minfi: a flexible and comprehensive Bioconductor package for the analysis of Infinium DNA methylation microarrays. Bioinformatics 30, 1363– 1369 (2014).

13. Zhou, W., Laird, P. W. & Shen, H. Comprehensive characterization, annotation and innovative use of Infinium DNA methylation BeadChip probes. Nucleic Acids Res 45, e22 (2017).

14. Maksimovic, J., Phipson, B. & Oshlack, A. A cross-package Bioconductor workflow for analysing methylation array data. F1000Res 5, 1281 (2016).

15. Ritchie, M. E. et al. limma powers differential expression analyses for RNA-sequencing and microarray studies. Nucleic Acids Res 43, e47 (2015).

16. Parekh, S., Ziegenhain, C., Vieth, B., Enard, W. & Hellmann, I. The impact of amplification on differential expression analyses by RNA-seq. Sci Rep 6, 25533 (2016).

17. Macosko, E. Z. et al. Highly Parallel Genome-wide Expression Profiling of Individual Cells Using Nanoliter Droplets. Cell 161, 1202–1214 (2015).

18. Leek, J. T., Johnson, W. E., Parker, H. S., Jaffe, A. E. & Storey, J. D. The sva package for removing batch effects and other unwanted variation in high-throughput experiments. Bioinformatics 28, 882–883 (2012).

19. Love, M. I., Huber, W. & Anders, S. Moderated estimation of fold change and dispersion for RNA-seq data with DESeq2. Genome Biol 15, 550 (2014).

20. Musa, J. & Cidre-Aranaz, F. Drug Screening by Resazurin Colorimetry in Ewing Sarcoma. Methods Mol Biol 2226, 159–166 (2021).

21. Wickham, H. et al. Welcome to the Tidyverse. Journal of Open Source Software 4, 1686 (2019).

22. Wickham, H. Reshaping Data with the reshape Package. Journal of Statistical Software 21, 1–20 (2007).

23. Dowle, M. et al. data.table: Extension of ‘data.frame’. (2023).

24. Getting Started with ggplot2 | SpringerLink. https://link.springer.com/chapter/10.1007/978-3-319-24277-4_2.

25. Jr, F. E. H. & functions), C. D. (contributed several functions and maintains latex. Hmisc: Harrell Miscellaneous. (2023).

26. Stein, C. K. et al. Removing batch effects from purified plasma cell gene expression microarrays with modified ComBat. BMC Bioinformatics 16, 63 (2015).

